# Morphological Identification of Ticks in district Swabi, Mardan, and Charsadda

**DOI:** 10.1101/2025.11.07.687134

**Authors:** Mohammad Adil, Hongsu He, Ahmad Ali, Jing Zhao, Yuefeng Wu, Uzair Alam, Junang Dai, Caihong Zhao, Tianyi Zhao, Hui Zhang, Zhihua Sun

## Abstract

Ticks are ectoparasitic arthropods that play a significant role in transmitting pathogens to humans and animals. This study aimed to address the research gap in understanding the morphological diversity and distribution of tick species in Swabi, Mardan, and Charsadda districts of Pakistan, where tick-borne diseases are prevalent but poorly documented. The primary objectives were to identify tick species, document their morphological characteristics, and map their distribution patterns across different habitats. Field surveys were conducted over nine months, collecting tick specimens from forests, grasslands, and residential areas. Specimens were preserved and analyzed using stereo-zoom microscopy and taxonomic keys for accurate classification. Key findings revealed a diverse tick fauna, including hard ticks (Ixodidae), with variations in body size, colour patterns, capitulum shape, and scutum features. Some species were widely distributed across all three districts, while others exhibited localized patterns. The study provides the first comprehensive morphological identification of tick species in these regions, highlighting their potential role in disease transmission. The implications of this research are significant for public health and disease control. The data can inform targeted interventions, enhance disease surveillance programs, and support the development of preventive measures to mitigate tick-borne diseases. By contributing to tick taxonomy and epidemiology, this study lays the groundwork for protecting public health in the study regions and beyond. This study addresses the lack of data on tick diversity in high-prevalence regions of Pakistan, providing critical insights for disease control and public health strategies. Limited information exists on the morphological diversity and distribution of tick species in Swabi, Mardan, and Charsadda districts. To identify tick species, document morphological characteristics, and map their distribution patterns in the study areas. Field surveys, specimen collection, stereo-zoom microscopy, and taxonomic classification were employed over nine months. Diverse tick fauna was identified, including hard ticks, with variations in morphology and distribution. Some species were widespread, while others were localized, highlighting potential disease transmission risks. The findings support targeted interventions, disease surveillance, and preventive measures, contributing to public health protection and tick-borne disease control in the region.

## 1. Introduction

Pakistan’s economy heavily relies on agriculture and livestock, with ticks posing a major threat to the latter. Ticks, hematophagous arthropod that feeds on the blood of their hosts, can damage livestock hides and reduce body weight (Ali et al., 2014). This parasitic infestation affects the economy by causing toxicosis, allergic reactions, and severe itching in livestock (Shah et al.). The prevalence of tick related disease is higher in tropical and subtropical regions (Muraleedharan, 2005). Given the economic impact of tick infestations, efforts are focused on controlling these parasites and the disease they transmit (Jongejan and Uilenberg. 2004).

Ticks first appeared in the Cretaceous period (146 to 66 million years ago) and continued to evolve and disperse until the Tertiary period (65 to 5 million years ago) (Funte de la et al., 2003). In Aristotle’s renowned book “Historia Animalium” dating back to 400 B.C., ticks were described as being produced from plants and as repulsive parasites. Throughout the mid-eighteenth century, parasitologists worldwide continued their studies on tick taxonomy, bionomy, prevalence, regional abundance, and seasonal patterns (Shah et al., 2015). The oldest known specimen of a tick, the Argasid bird tick, was found in Cretaceous New Jersey amber, while more recent specimens from Dominican and Baltic ambers can be classified into existing taxa (Dunlop et al., 2016). Some tick species discovered in preserved reptiles suggest an origin dating back around 99 million years, such as the Deinocroton Dracula tick found in a dinosaur feather (Penalver et al., 2017).

Ticks are common parasites found both in both humans and animals worldwide, especially in tropical and subtropical regions. They are significant disease vectors, spreading over 200 diseases through around 10% of the 899 tick species. Ticks not only suck blood but also have other direct negative effects on their hosts (Jongejan and Uilenberg., 2004).

Ticks play a significant role in the global spread of tick-borne animal diseases, serving as important vectors for various illnesses. In Pakistan, ticks, as ectoparasites, transmit different protozoans that effect the growth rate of domestic animals, contaminate animal products, and contribute the spread of diseases such as theileriosis and babesiosis (Ghosh et al., 2007; Lubud and Despommier et al. 2000; Jongejan and Uilenberg 2004). Additionally, tick can transmit zoonotic agents including bacteria (*Francisella tularensis, Borrelia burgdorferi*), rickettsia (*Rickettsia rickettsii, Coxiella burnetii, Ehrlichia canis*), and protozoa (*Babesia divergens, B. microti*), posing new challenges to the healthcare industry due to the increasing number of cases worldwide (Kilpatrick & Randolph, 2012). The global rate of urbanization has increased in recent decades, leading to significant changes in the diversity of flora and fauna. This has impacted the transmission of human and animal diseases by vectors and altered the interactions between animals and pathogens. (Bradley and Altizer, 2007; Decker et al., 2010). Ticks, ectoparasites are found on animals, is the second-largest category of transmission vectors for zoonoses, capable of spreading various infections including bacteria, viruses, and parasites (Boulanger et al., 2019). Tick bites can harm the host’s reproductive capacity, leading to anemia, emaciation, skin damage, inflammation, and rashes. In severe cases; tick bites can cause paralysis and even death (Fourie et al., 2013).

Many investigators have carried out researches on different parameters of ticks in different regions in Pakistan but no morphological identification of ticks from district Swabi, Mardan and Charsadda have been described before so, there is a demand for the appropriate morphological identification of ticks from these regions.

In the current study, morphology of various species from different fauna were studied in detail, certain tick species with known zoonotic potential were also identified, highlighting the public health significance of this research. Understanding the morphological diversity and distribution of ticks in the region is essential for devising effective control and management strategies to mitigate tick-borne diseases’ impact on human and animal health.

## 2. Methods and Materials

### 2.1 Study Design

The cross-sectional study was conducted between June 2022 and March 2023. Tick samples were collected from domesticated animals (such as cows, dogs, buffalo, goats, sheep and camels during routine visits to farms and grazing grounds in the research regions. A predetermined proforma was used to document the animal’s species, age, location, and sex. Various body parts of the animals, including shoulders, dewlaps, bellies, heads, ears, necks, backs, legs, perineum, and tails, were examined for ticks. Ticks were carefully removed from the animals’ skin using forceps and preserved in Eppendorf tubes for transportation to the parasitology lab. The collected specimens were cleaned with distilled water and stored in a separate container with 70% ethanol. Using a stereomicroscope with 100–200-fold magnification and a digital microscope, adult ticks (both male and female) were morphologically identified. Common taxonomic keys were used for classification; they were categorized based on their morphological traits according to Walker’s classification.

### 2.2 Collection Sites for Sample

Ticks were collected from various areas in Charsadda, Mardan, and Swabi. In Charsadda, the collection sites included Umarzai, Turangzai, Utmanzai, Mera Turangzai, Dedarabad Turangzai, Esazai, Badra Khiel, Banda Turangzai, Rajjar, Ameer Abad, Mandani, Merozai Turangzai, Peran Turangzai and Malak Abad.

In Mardan, the collection sites were Rustam, Ale Landi, Daggar, Bhai khan, Landai, Gujrat, Bakhshali, Maqam Chowk, Pakistan Chowk, Toru, Mayar, Shahbaz Garhi, Hussai, and Garyala.

In Swabi, the collection sites included were Upper Maniray, Lower Maniray, Gulu Deray, Steppa Neer, Shwe Kali, Nawi Kali, Tolanday, Permolu, Hisha Marali, Sherdara, Dagai, Tarakay, Yar Husain, Dobian, Babu Derai and Khas Bazargi (Figure. 1).

**Figure 1.**
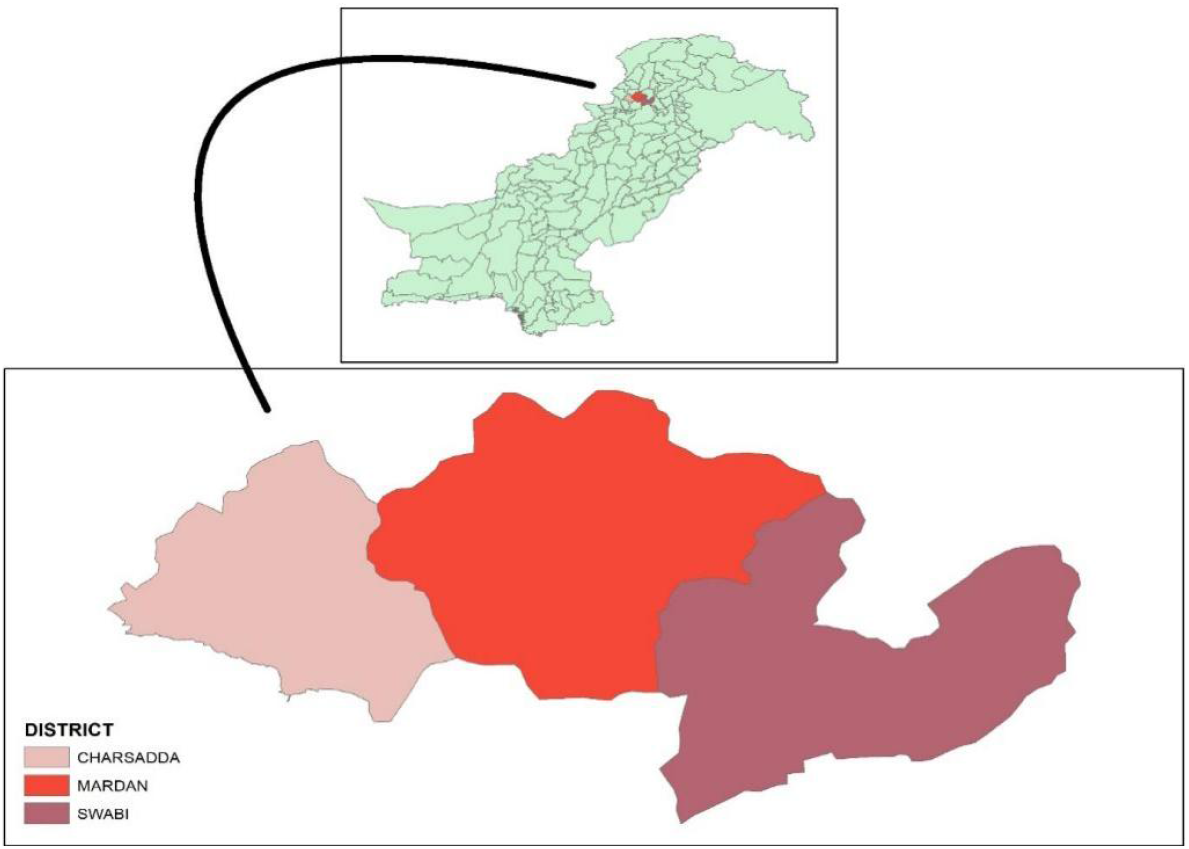
Shows the study map for district Charsadda, Mardan and Swabi.

### 2.3 Target Hosts

The target hosts in three districts, Charsadda, Mardan, and Swabi, included goats, camels, dogs, donkeys, cows, sheep, bulls, and calves. Our focus was on soft ticks, but unfortunately, the ticks were not found on the host.

Physical host control without chemical injection was carried out on domestic animals during tick isolation and search on their body surfaces.

### 2.4 Tabulation

The table was created using Excel and includes the tube number, target host’s common name, scientific name, host gender, and location. It also includes the collection date and meteorological data such as temperature, humidity, and precipitation. The number of ticks removed from the host, their geographic coordinates, tick species, and life stage were recorded after identification.

### 2.5 Tick identification

To identify the tick genus, the dorsal portion of the tick was affixed to a microscope for examination. The primary characteristic examined was the mouth parts followed by the lateral groove, festoon, and Scutum form on the dorsal side for species identification of males and females. The ventral side of the tick was then observed under a microscope to check the ventral plate of the males for species determination, along with investigations on the lateral side’s spiracle plate. The morphology of the vaginal opening was investigated in both nymphs and adults to aid in species identification. Furthermore, the leg color and leg number were studied to identify species and determine developmental phases.

### 2.6 *Hyalomma* genus Morphological identification

In contrast to other genera, *Hyalomma* ticks are medium sized and unfed, measuring 5 to 6 millimeters in length, including the mouth. They have a medium angular lateral border on the basis capituli, long frontal teeth, three paired ventral plates in males, and equal paired spurs on the coxae. Their legs have rings of a light color, with Articles 1 and 3 of the Palp shorter than Article 2. The scutum and scutum color are brown, and they have convex eyes. Both males and females have a festoon, are not feed, and exhibit an anal groove and a sizable spiracle plate posterior to leg 4.

### 2.7 *Rhipicephalus* genus Morphological identification

The *Rhipicephalus* genus is characterized by medium-sized, unfed ticks measuring 3 to 5 millimeters, including the mouthpart. They have an anterior mouthpart with tiny palp articles, a hexagonal basis capituli with angular lateral borders, no rings of light color on the legs, and a dark black scutum in males and conscutum in females. The eyes can be flat or somewhat convex, males have festoons, and females have a large posterior spiracle plate on leg 4. Males have two paired ventral plates, uneven spurs on coxae 1, and a groove anterior to the anus.

### 2.8 *Haemaphysalis* genus Morphological identification

The *Haemaphysalis* genus is distinct from others due to its medium sized unfed ticks measuring 3 millimeters, including the mouthpart, and lacking eyes. The basis capituli has a rectangular shape with straight lateral borders and short anterior mouth-parts. In some species, palp 2 is wide and has a characteristic shape. These ticks have dark-colored legs, with males having a scutum and females having conscutum. Both males and unfed females exhibit festoons, and the posterior large spiracle plate is present on leg 4. Males do not have ventral plates, and Coxae 1 has a single spur.

## 3. Results

### 3.1 The distribution of ticks from different host in various regions

In the entire sample collected from Jun 2022 to March 2023, a total of 89 animals were observed, including cows (870 ticks), dog (67 ticks), sheep and goats (534 ticks), camels (47 ticks), and buffaloes (304 ticks), totaling 1822 ticks.

In district Charsadda, a total of 530 ticks were collected from 132 hosts. The distribution of tick species was as follows: *R. microplus* accounted for 41.50% (220 ticks), *R. haemaphysaloides* for 4.5% (24 ticks), *R. sanguineus* for 10.3% (55 ticks), *R. turanicus* for 18.3% (97 ticks), *Ha. Bispenosa* for 0% (0 ticks), *Hy. Anatolicum* for 18.7% (100 ticks), *Hy. Dromedarii* for 11.88% (63 ticks), and *Hy. Scupense* for 4.1% (22 ticks) (Table 1). The P-Value of 0.05 was calculated using Chi square test, indicating a correlation between the observed frequencies of the tick species and their distribution (Figure 2). This suggests that the distribution of tick species is not random, indicating that certain species may be more prevalent than others.

**Table 1.** shows tick prevalence in Charsadda.

| S.No | Name of tick species | Number of total ticks | % |
| --- | --- | --- | --- |
| 1 | *R. Microplus* | 220 | 41.50% |
| 2 | *Hy. Dromedarii* | 10 | 11.88% |
| 3 | *R. Haemaphysaloides* | 24 | 4.5% |
| 4 | *R. Sanguineous* | 55 | 10.3% |
| 5 | *R. Turanicus* | 97 | 18.3% |
| 6 | *Hy. Anatolicum* | 100 | 18.7% |
| 7 | *Hae. Bispenosa* | 0 | 0% |
| 8 | *Hy. Kumari* | 1 | 0.18% |
| 9 | *Hy. Scupense* | 22 | 4.1% |
| 10 | *Hae. Montgomeryi* | 1 | 0.18% |
|  | Total | 530 | 100% |

**Figure 2.**
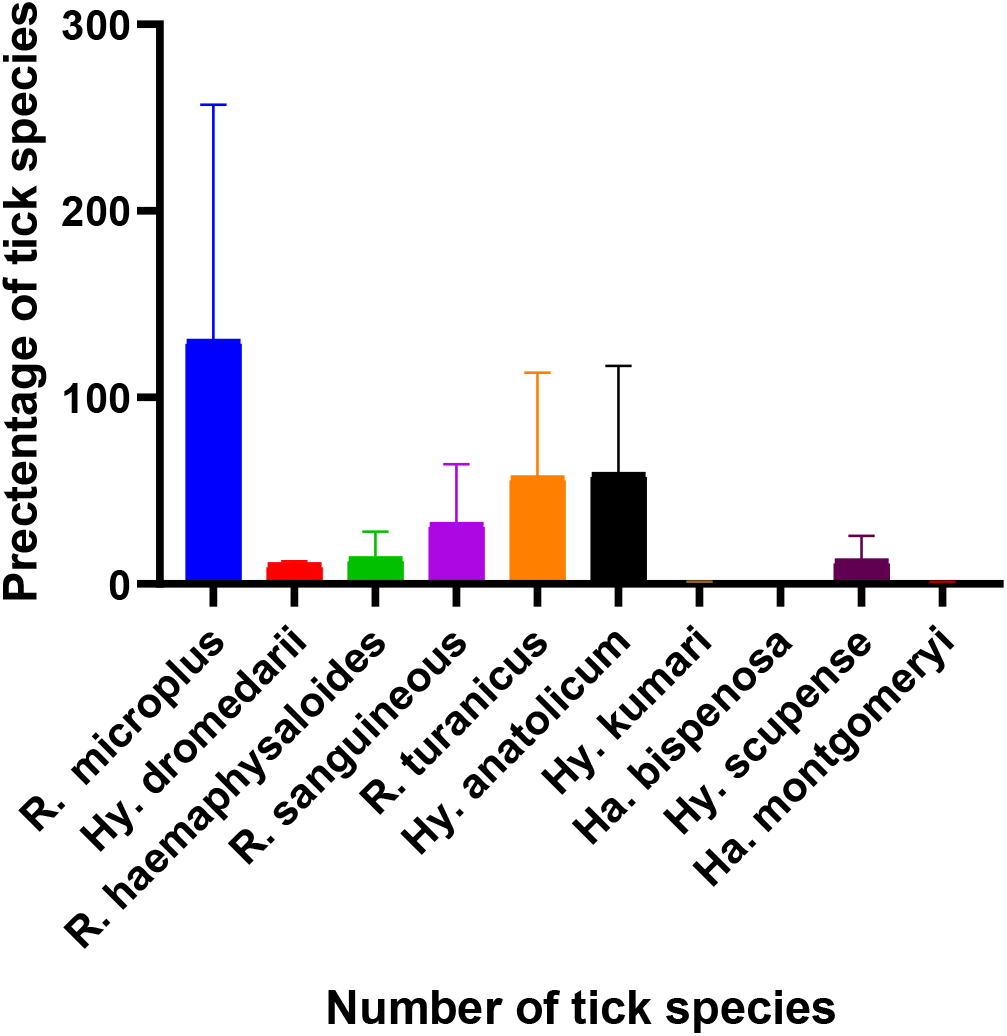
Shows number of spp. with percentages in district Charsadda.

A total of 571 ticks were collected from 121 hosts in district Mardan. The distribution of tick species was as follow: *R. microplus* accounted for 26.26% (150 ticks), *R. haemaphysaloides* for 4.20% (24 ticks*), R. sanguineous* for 16.63% (95 ticks), *R. turanicus* for 26.26% (150 ticks), *Hae. Bispenosa* for 0% (0 ticks), *Hy. Anatolicum* for 26.26% (120 ticks), *Hy. Dromedary* for 1.70% (10 ticks), *Hy. Scupense* for 1.92% (21 ticks) and *Hae. Montgomeryi* for 0.17% (11 ticks) (Table.1) (Figure 2).

A total of 571 ticks were collected from 121 hosts in district Mardan. The distribution of tick species was as follows: *R. micro plus* accounted for 26.26% (150 ticks), *R. haemaphysaloides* for 4.20% (24 ticks), *R. sanguineous* for 16.63% (95 ticks), *R. turanicus* for 26.26% (150 ticks), *Ha. Bispenosa* for 0% (0 ticks), *Hy. Anatolicum* for 26.26% (120 ticks), *H. dromedary* for 1.75% (10 ticks), *Hy. Scupense* for 1.92% (21 ticks), and *Ha. Montgomeryi* for *0.17%* (11 ticks). (Figure 3) (Table 2).

**Table 2.**
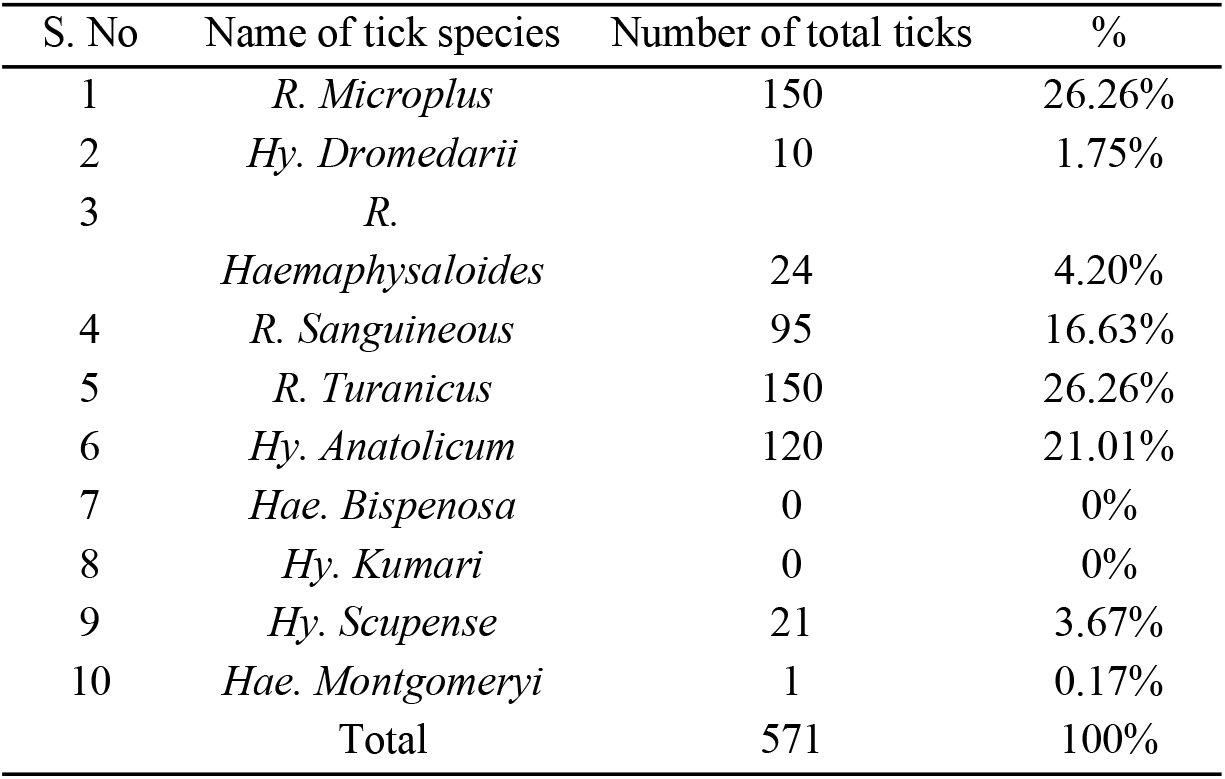
Show tick prevalence in district Mardan.

| S. No | Name of tick species | Number of total ticks | % |
| --- | --- | --- | --- |
| 1 | *R. Microplus* | 150 | 26.26% |
| 2 | *Hy. Dromedarii* | 10 | 1.75% |
| 3 | *R.*<br>*Haemaphysaloides* | 24 | 4.20% |
| 4 | *R. Sanguineous* | 95 | 16.63% |
| 5 | *R. Turanicus* | 150 | 26.26% |
| 6 | *Hy. Anatolicum* | 120 | 21.01% |
| 7 | *Hae. Bispenosa* | 0 | 0% |
| 8 | *Hy. Kumari* | 0 | 0% |
| 9 | *Hy. Scupense* | 21 | 3.67% |
| 10 | *Hae. Montgomeryi* | 1 | 0.17% |
|  | Total | 571 | 100% |

**Figure 3.**
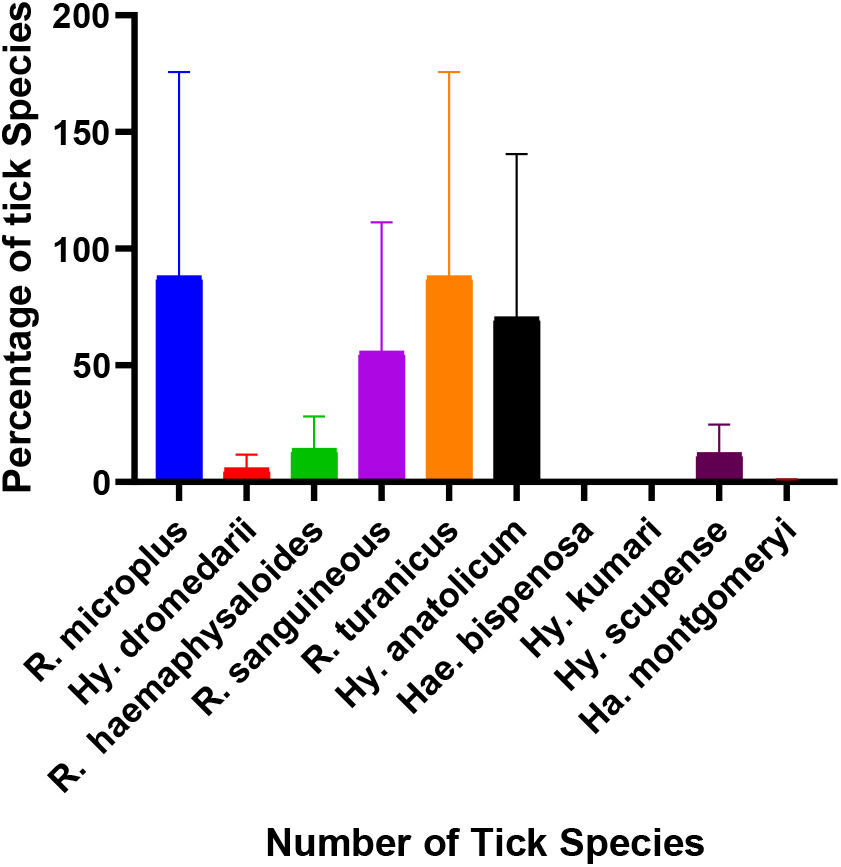
Shows number of spp. with percentages in district Mardan.

A total of 721 ticks were collected from 197 hosts in the Swabi district. Among them, *R. micro plus* accounted for 26.26% (330 ticks), *R. haemaphysaloides* for 3.74% (27 ticks), *R. sanguineous* for 13.86% (100 ticks), *R. turanicus* for 23.57% (170 ticks), *Ha. Bispenosa* for 0.27% (2 ticks), *Hy. Anatolicum* for 4.85% (35 ticks), *Hy. Dromedarii* for 3.46% (25 ticks), *Hy. Scupense* for 1.80% (26 ticks).

In the entire sample collected from Jun 2022 to March 2023, a total of 89 animals were observed, including cows (870 ticks), dog (67 ticks), sheep and goats (534 ticks), camels (47 ticks), and buffaloes (304 ticks), totaling 1822 ticks. (Table 3) (Figure 4).

**Table 3.**
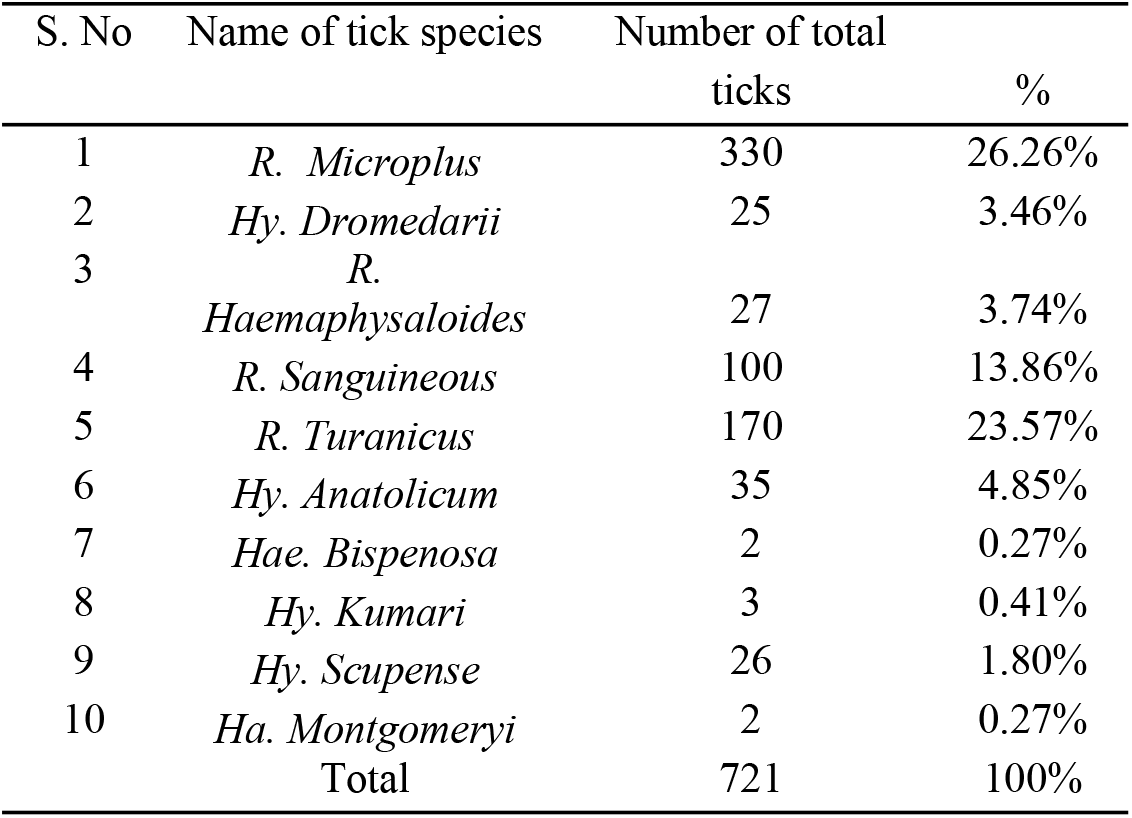
Shows tick prevalence in district Swabi.

| S. No | Name of tick species | Number of total<br>ticks | % |
| --- | --- | --- | --- |
| 1 | *R. Microplus* | 330 | 26.26% |
| 2 | *Hy. Dromedarii* | 25 | 3.46% |
| 3 | *R. Haemaphysaloides* | 27 | 3.74% |
| 4 | *R. Sanguineous* | 100 | 13.86% |
| 5 | *R. Turanicus* | 170 | 23.57% |
| 6 | *Hy. Anatolicum* | 35 | 4.85% |
| 7 | *Hae. Bispenosa* | 2 | 0.27% |
| 8 | *Hy. Kumari* | 3 | 0.41% |
| 9 | *Hy. Scupense* | 26 | 1.80% |
| 10 | *Ha. Montgomeryi* | 2 | 0.27% |
|  | Total | 721 | 100% |

**Figure 4.**
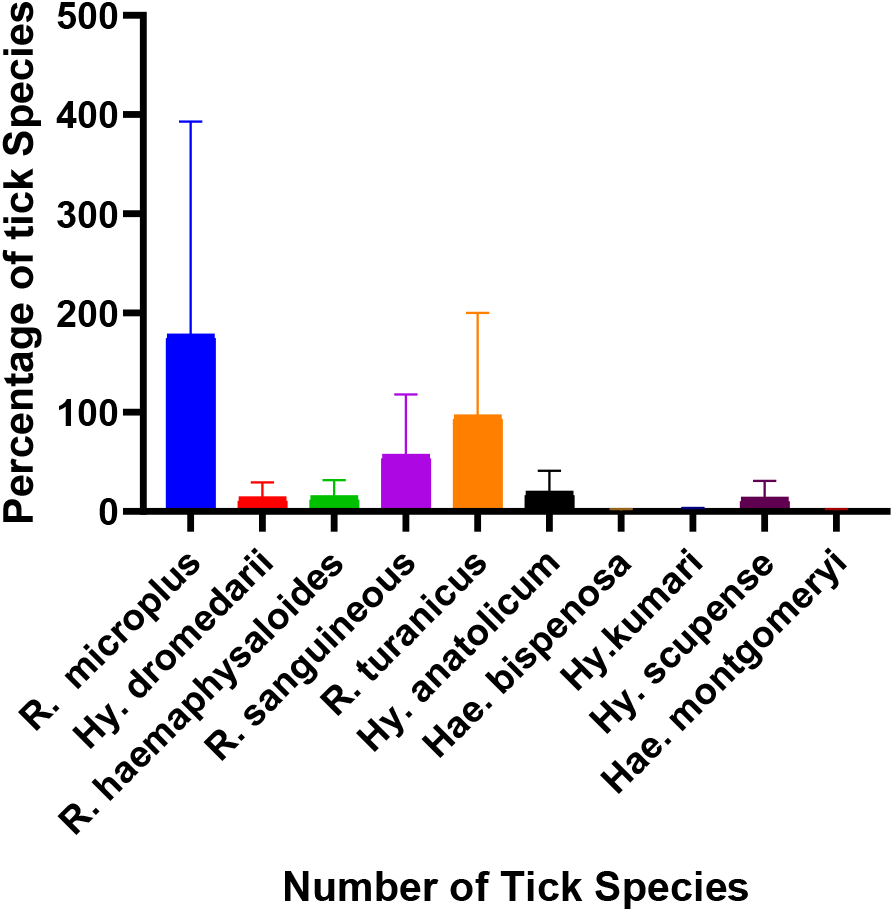
Shows number of spp. with percentages in district Swabi.

### 3.2 Morphological identification of ticks by microscope

*Rhipicephalus microplus*, commonly known as the southern cattle tick, is a one-host tick native to Asia but has been introduced to various regions worldwide, such as Africa, Australia, North and South America, and Europe. It is a significant pest of cattle and other livestock and can transmit diseases babesiosis, anaplasmosis, and theileriosis. Controlling *R. Microplus* is challenging due to its resistance to numerous acaricides. *Rhipicephalus microplus* differs from other common tick species in its host range, life cycle, geographical distribution, and the diseases it transmits. It is important to be aware of the tick species present in a specific area to take necessary precautions to prevent tick bites and avoid tick-borne diseases (Figure 5).

**Figure 5.**
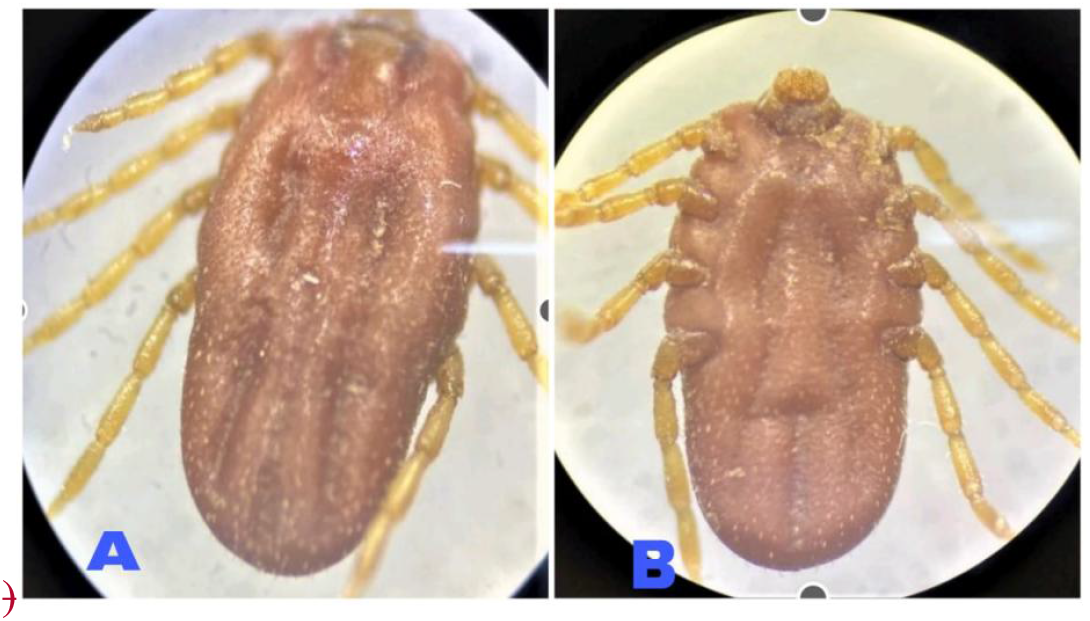
Shows the *Rhipicephalus microplus* female collected from a cow. A, *Rhipicephalus microplus* female dorsal view. B, *Rhipicephalus microplus* female ventral view.

Adult *Rhipicephalus sanguineous* ticks relatively small compared to other tick species, with a color range from dark brown to reddish brown. The female tick body swells more after feeding; whereas the male tick body is typically longer and flatter from top to bottom. Due to its excellent adaptability to dogs, *Rhipicephalus sanguineous* commonly infest them, especially in places like kennels and dogs’ shelters where dogs are kept in close quarters. Ticks are often removed directly from dog’s skin during veterinary visits. Dogs and human can contract various infections from *Rhipicephalus sanguineous*, including ehrlichiosis and babesiosis which seriously impact a dog’s health (Figure 6).

**Figure 6.**
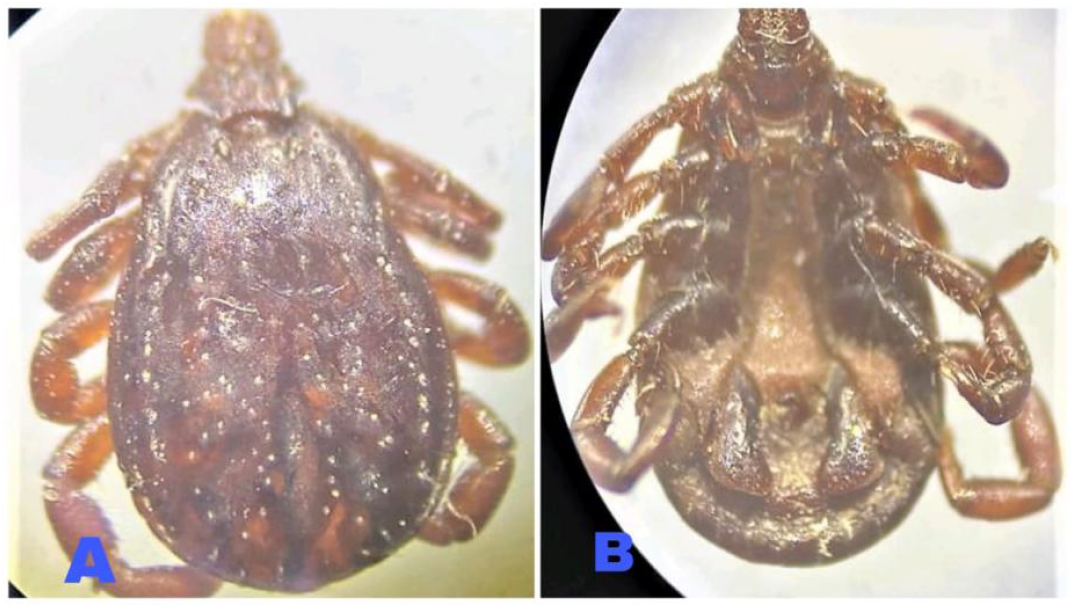
Shows the *Rhipicephalus sanguineus* male collected from a dog. A, *Rhipicephalus sanguineus* male dorsal view. B, *Rhipicephalus sanguineus* male ventral view.

Similar to other ticks, the male *Rhipicephalus haemaphysaloides* can be identified by its distinctive features. It has a scutum or shield on the dorsal side that covers part of its body. The mouth parts, including the chelicerae and hypostome, are visible on the ventral side and are adapted for piercing the host skin to feed on blood (Figure 7).

**Figure 7.**
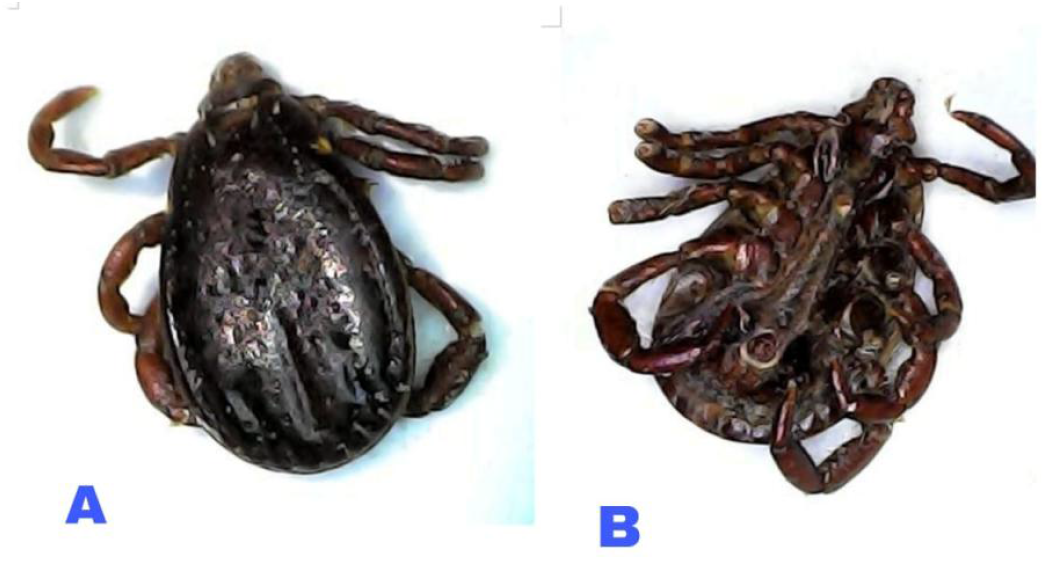
*R. haemaphysaloides* male adult collected from cattle. A, *R. haemaphysaloides* male adult dorsal view. B, *R. haemaphysaloides* male adult ventral view.

Female *Rhipicephalus turanicus* ticks play a vital role in the ecosystem by regulating host populations and serving as a host for various parasites and diseases. However, they also pose a threat as potential vectors of infections to human and animals, emphasizing the importance of research and management efforts for public health and veterinary purposes. Studying female *Rhipicephalus turanicus* ticks collected from sheep can offer valuable understandings into their behavior, biology, and role in disease transmission (Figure 8).

**Figure 8.**
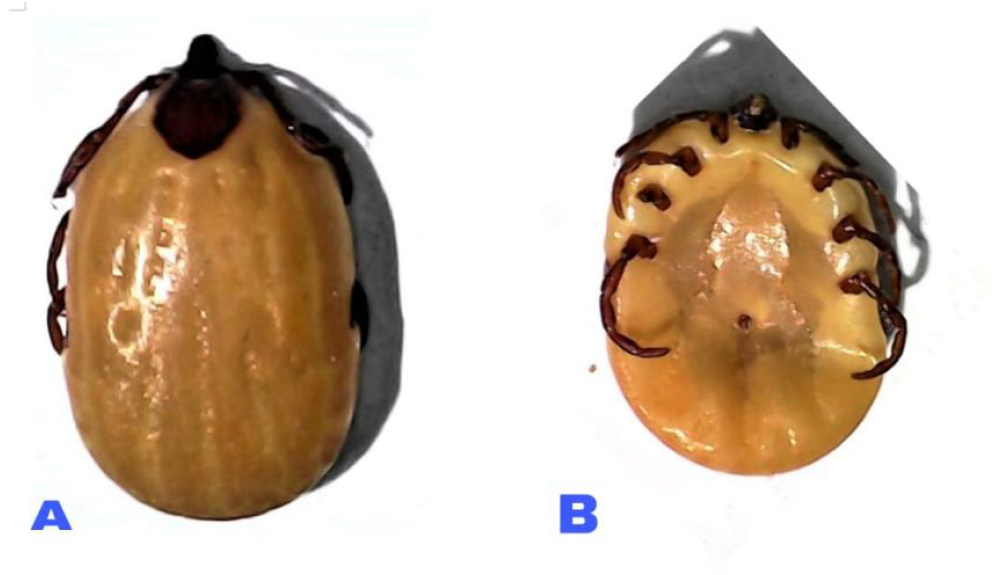
Shows the *Rhipicephalus turanicus* female collected from sheep. A, *Rhipicephalus turanicus* female dorsal view. B, *Rhipicephalus turanicus* female ventral view.

*Hyalomma anatolicum* ticks are commonly found infecting cattle, among other hosts such as sheep, goats, and wild animals. Compared to other tick species, the female *Hyalomma anatolicum* tick is relatively large, measuring several millimeters in length. It has a hard sclerotized exoskeleton typically of hard ticks, covering an elongated, flattened dorsoventrally body. The body color ranges from brown to grey before feeding, but darkens after meal (Figure 9).

**Figure 9.**
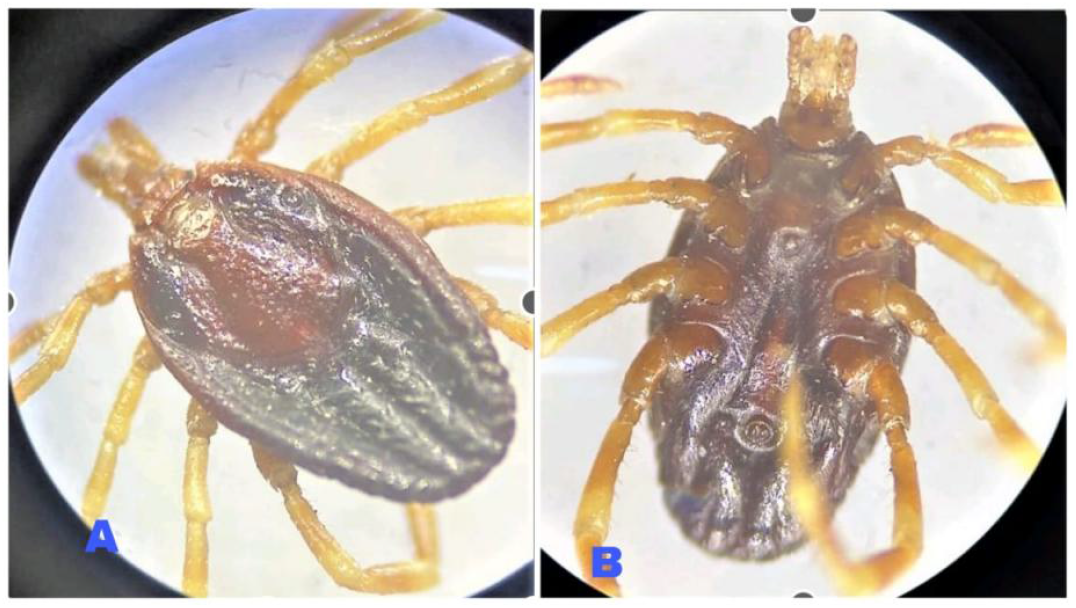
Show *Hyalomma anatolicum* female collected from cattle. A, *Hyalomma anatolicum* male female dorsal view. B, *Hyalomma anatolicum* female adult ventral view.

Female *Hyalomma Scupense* ticks can grow up to 15 millimeters in length when fully engorged, larger than males. Unfed females typically measure between 4 to 6 millimeters. They exhibit a range of colors typically ranging from dark brown to reddish brown (Figure 10).

**Figure 10.**
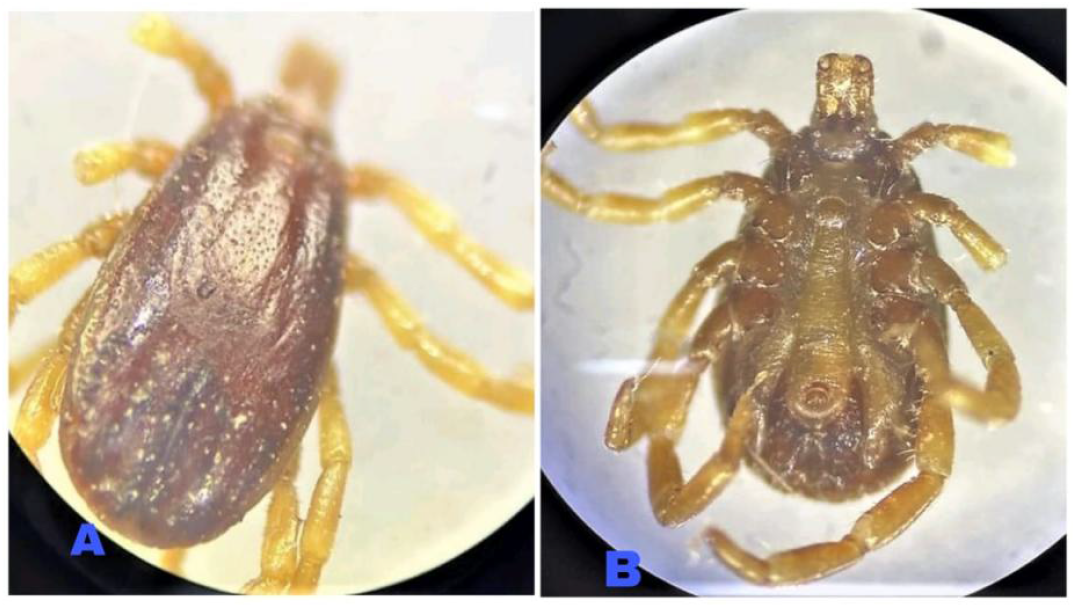
*Hyalomma Scupense* female adult collected from sheep. A, *Hyalomma Scupense* female adult dorsal view. B, *Hyalomma Scupense* female adult ventral view.

Female *Hyalomma kumari* female ticks play an important role in the ecosystem by helping regulate host populations and acting as hosts for various parasites and pathogens. However, they also present a significant risk as potential vectors for diseases that affect both humans and animals. This highlights the need for ongoing research and effective management strategies to protect public health and animal well-being. Studying female *Hyalomma kumari* ticks, especially those collected from goats, provides valuable insights into their behavior, biology, and their role in the transmission of infections (Figure 11).

**Figure 11.**
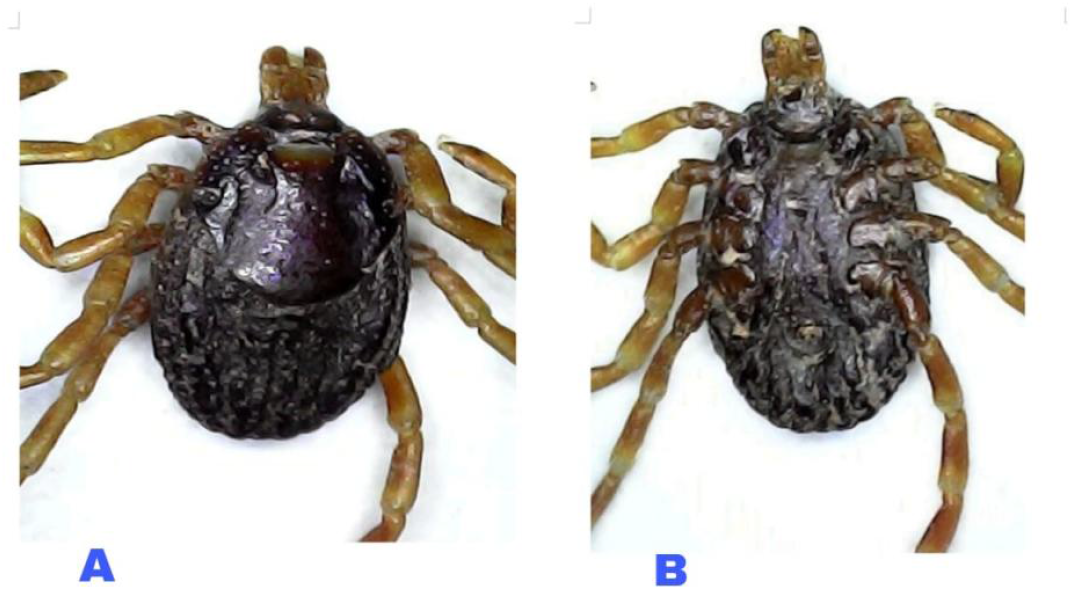
*Hyalomma kumari* female adult collected from goat. A, *Hyalomma kumari* female adult dorsal view. B, *Hyalomma kumari* female adult ventral view.

Male *Hyalomma dromedarii* adult ticks, collected from camels, play a vital role in the ecosystem by influencing host populations and acting as vectors for various parasites and pathogens. These ticks are particularly concerning because they are known to transmit several diseases, including Crimean-Congo hemorrhagic fever (CCHF), Theileriosis, and Babesiosis. These diseases can have severe consequences for both human and animal health. As *Hyalomma dromedarii* ticks feed on their hosts, they facilitate the spread of these pathogens, making them significant vectors in disease transmission. This underscores the critical need for targeted research and management strategies to mitigate the risks associated with these ticks. Studying male *Hyalomma dromedarii* adult ticks, especially those collected from camels, provides essential insights into their behavior, biology, and their role in the epidemiology of tick-borne diseases (Figure 12).

**Figure 12.**
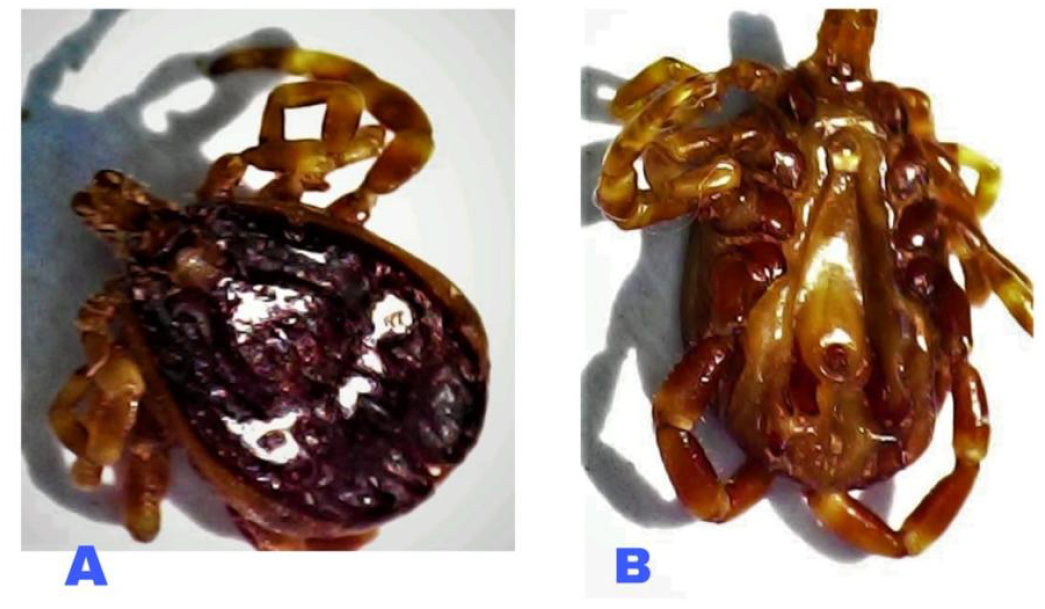
*Hyalomma dromedarii* male adult collected from camel. A, *Hyalomma dromedarii* male adult dorsal view. B, *Hyalomma dromedarii* male adult ventral view

Male *Hyalomma kumari* ticks play an essential role in the ecosystem by helping regulate host populations and serving as hosts for various parasites and pathogens. However, they also pose a risk as potential vectors for diseases that can affect both humans and animals. This underscores the need for continued research and management efforts to safeguard public health and animal welfare. Studying male *Hyalomma kumari* ticks, especially those collected from goats, provides valuable insights into their behavior, biology, and their contribution to the transmission of infections (Figure 13).

**Figure 13.**
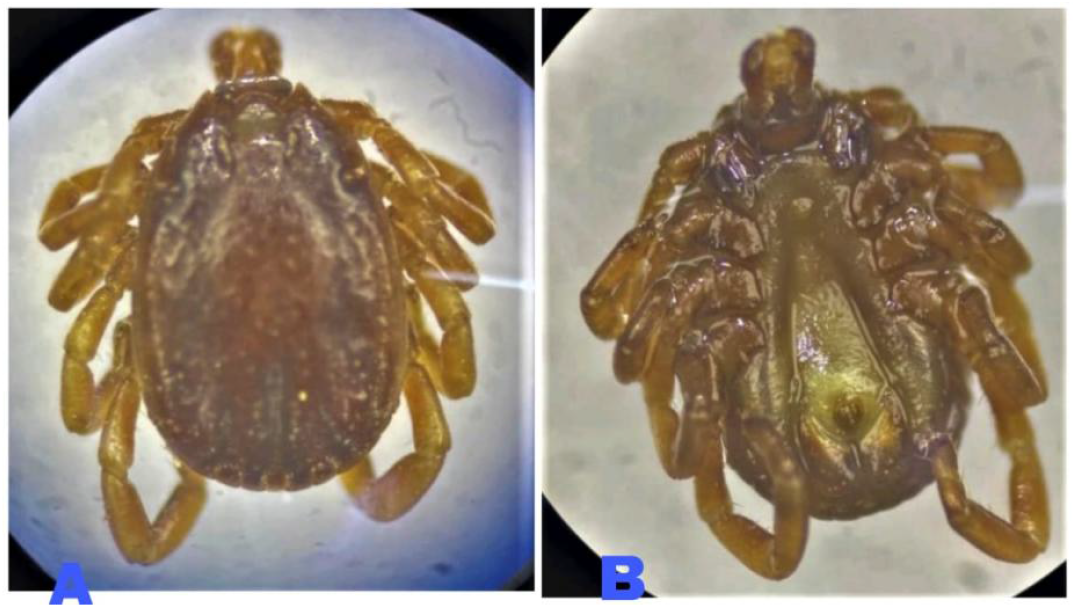
*Hyalomma kumari* male adult collected from goat. A, *Hyalomma kumari* male adult dorsal view. B, *Hyalomma kumari* male adult ventral view.

Male *Haemaphysalis Montgomery* adult ticks, collected from goats, play a significant ecological role by regulating host populations and serving as vectors for various pathogens. These ticks are known to transmit diseases such as Tick-borne encephalitis (TBE) and Anaplasmosis, which can affect both livestock and humans. As they feed on their hosts, they facilitate the spread of these pathogens, posing a serious threat to animal health and public safety. Given the potential for disease transmission, studying male *Haemaphysalis Montgomery* adult ticks, particularly those collected from goats, is essential for understanding their biology, behavior, and role in the transmission of tick-borne diseases. This knowledge is crucial for developing effective control and prevention measures to mitigate the impact of these ticks on livestock and human health (Figure 14).

**Figure 14.**
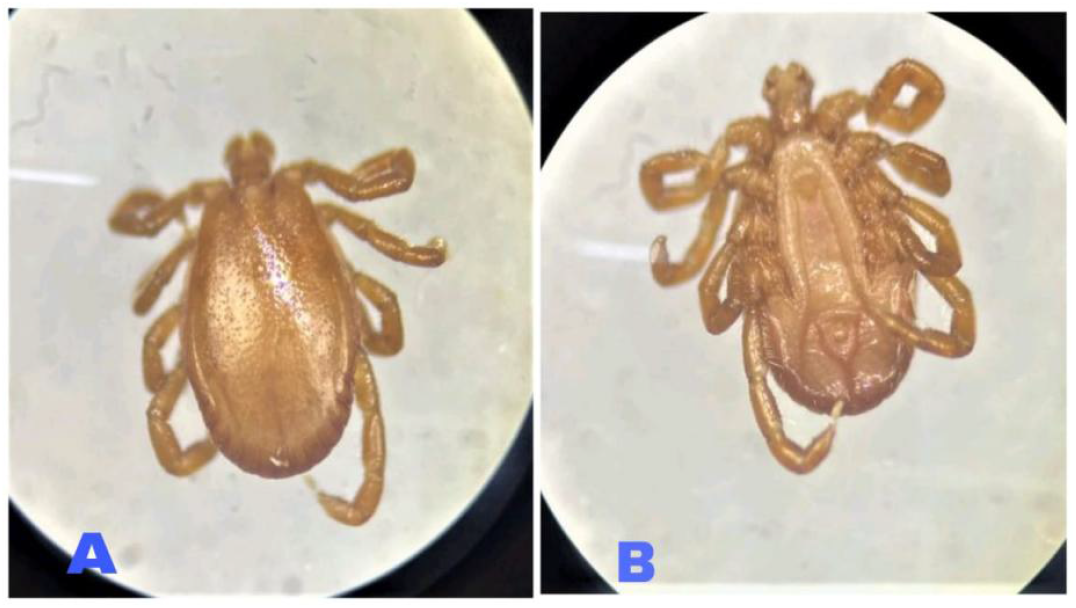
Show the *Haemaphysalis Montgomery* male adult collected from goat. A, *Haemaphysalis Montgomery* male adult dorsal view. B, *Haemaphysalis Montgomery* male adult ventral view.

Female *Haemaphysalis Bispenosa* adult ticks, collected from goats, serve as both ecological regulators and vectors of significant pathogens. These ticks are primarily involved in the transmission of protozoan parasites such as Babesia spp. and Theileria spp., which are causative agents of Babesiosis and Theileriosis, respectively. These diseases have a profound impact on livestock, leading to significant morbidity and economic loss, particularly in endemic regions. *Haemaphysalis Bispenosa* ticks acquire these pathogens during their blood meals from infected hosts and subsequently transmit them during subsequent feeding events. Understanding the biology, feeding behavior, and vector competence of *Haemaphysalis Bispenosa*, particularly in the context of their interaction with goats, is essential for elucidating their role in the epidemiology of tick-borne diseases. Detailed studies on the population dynamics, life cycle, and pathogen transmission efficiency of these ticks are crucial for developing targeted control measures aimed at reducing the incidence of tick-borne diseases in livestock populations (Figure 15).

**Figure 15.**
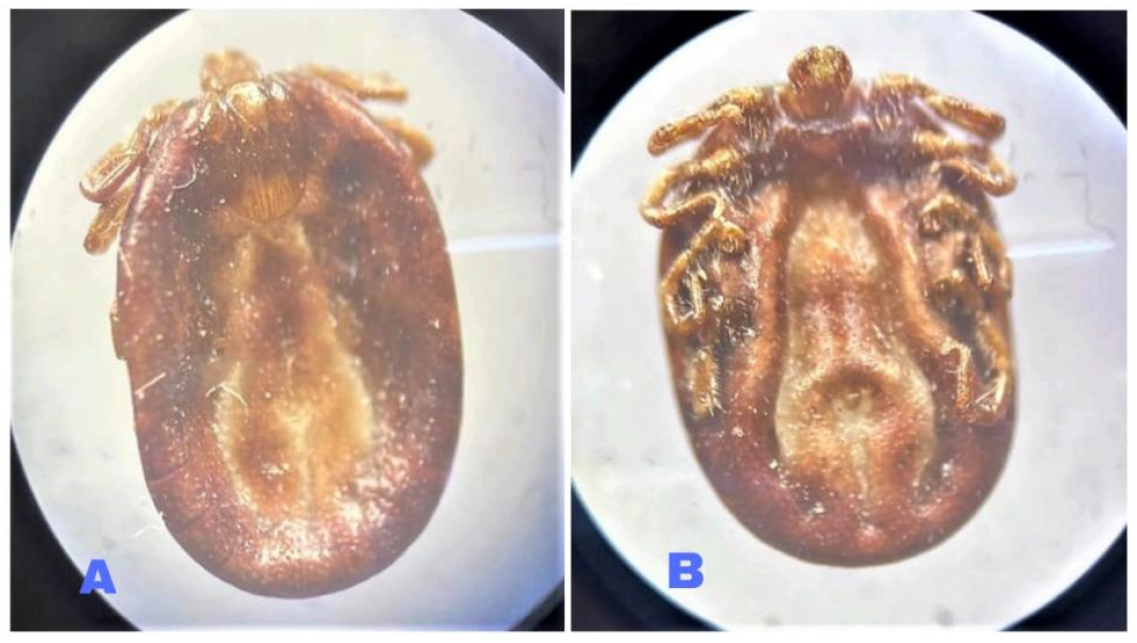
Shows the *Haemaphysalis bispenosa* female adult collected from goat. A, *Haemaphysalis bispenosa* female adult dorsal view. B, *Haemaphysalis bispenosa* female adult ventral view.

## 4. Discussion

In Pakistan, studies have revealed that ticks are rare in terms of their morpho-taxonomy, distribution, prevalence, molecular phylogeny, and management methods (appropriate immunization), as well as the research region. The purpose of the study, research questions, and literature review, and synthesizes the findings to draw meaningful conclusions. It also offers recommendations for practice and future research to address the gaps identified in the study. The purpose of this study was to investigate the morpho-taxonomy, distribution, prevalence, and molecular phylogeny of ticks in Pakistan, with a focus on their impact on livestock and the associated economic losses. The research questions centered on identifying the predominant tick species, their seasonal distribution, predilection sites on hosts, and the factors influencing their prevalence. The literature review highlighted the limited understanding of tick taxonomy in Pakistan, the economic burden of tick-borne diseases, and the lack of molecular-level verification of tick species. The methodology involved field surveys, morphological identification, and molecular analysis of ticks collected from cattle in Swabi, Mardan, and Charsadda. The findings revealed the presence of three major tick genera—*Hyalomma, Rhipicephalus*, and *Haemaphysalis*—with *Rhipicephalus microplus* being the most prevalent species. Seasonal trends showed higher tick infestation during the summer months, particularly in July, due to favourable environmental conditions such as high temperature and humidity. The study identified *Hyalomma* (e.g., *Hy. anatolicum, Hy. kumari, Hy. dromedarii*, and *Hy. Scupense*) and *Rhipicephalus* (e.g., *R. microplus, R. sanguineus*, and *R. turanicus*) as the dominant tick genera in the study area. These findings align with previous studies by Karim et al. (2017) and Jamil et al. (2022), who also reported the prevalence of these genera in Pakistan. The high prevalence of *R. microplus* (26.26%) and *R. turanicus* (23.57%) underscores their significance as vectors of tick-borne diseases such as bovine tropical theileriosis and babesiosis. The morphological complexity of the *Boophilus* subgenus, as noted by Barker and Walker (2014), further complicates their identification and management. The study confirmed that tick infestation peaks during the summer months (June to September), with the highest prevalence in July. This finding is consistent with the work of Ali et al. (2019) and Islam et al. (2006), who also reported higher tick activity during warm and humid conditions. The low infestation rates in winter (November) highlight the influence of environmental factors such as temperature and humidity on tick populations. These seasonal trends are critical for developing targeted control strategies, such as timed acaricides applications and vaccination programs. The face and ears of cattle were identified as the most common predilection sites for ticks, with a prevalence of 62%. This finding corroborates the work of Rafique et al. (2015), who also noted similar predilection sites. The study also found that adult cattle were more susceptible to tick infestation than young cattle, which aligns with the observations of Mattioli et al. (1998). Host factors such as age, gender, and breed, along with environmental variables like vegetation and geographical features, significantly influence tick distribution and abundance. The high prevalence of tick-borne pathogens such as *Babesia bovis, Babesia bigemina*, and *Anaplasma marginale* poses a significant threat to livestock health and productivity. The economic losses associated with these diseases, as highlighted by Jabbar et al. (2015), underscore the need for effective tick management strategies. The lack of molecular-level verification of tick species, as noted by Karim et al. (2017), further complicates disease control efforts.

To address these challenges, several recommendations for practice are proposed. First, improved tick identification and taxonomy are essential. Developing and disseminating morphological and molecular keys for accurate tick identification can help address the taxonomic discrepancies highlighted in this study. Second, seasonal control strategies should be implemented, such as targeted acaricides applications and vaccination programs during peak tick activity periods (June to September) to reduce infestation rates. Third, farmer education is critical. Educating farmers on the importance of regular tick inspections, proper acaricides use, and the role of environmental management in controlling tick populations can significantly reduce the burden of tick-borne diseases. Finally, integrated pest management (IPM) practices should be promoted. Combining chemical, biological, and cultural control methods can reduce reliance on acaricides and minimize resistance development.

For future research, several areas warrant further investigation. First, molecular phylogeny studies should be conducted to verify the identity of tick species and explore their phylogenetic relationships. Second, pathogen surveillance is needed to investigate the prevalence and distribution of tick-borne pathogens in different regions of Pakistan to identify high-risk areas. Third, host-pathogen interactions should be studied to better understand disease transmission dynamics. Fourth, the impact of climate change on tick distribution and abundance should be assessed to predict future trends and develop adaptive management strategies. Finally, economic impact studies should quantify the economic losses caused by tick-borne diseases to justify investments in control and prevention programs.

This study provides valuable insights into the morpho-taxonomy, distribution, and prevalence of ticks in Pakistan, highlighting the need for improved identification methods, targeted control strategies, and further research to mitigate the economic and health impacts of tick-borne diseases.

## 5. Conclusion

In conclusion, the study highlights the significant issue of tick infestations in the districts of Mardan, Swabi, and Charsadda, where ticks of the genera *Rhipicephalus, Hyalomma*, and *Haemaphysalis* are prevalent across various animal hosts. The key findings reveal that *R. microplus* and *R. sanguineus* are the most frequently infesting species, with *Hyalomma* species (*Hy. dromedarii, Hy. anatolicum*, and *Hy. suspense*) also posing a considerable threat. The geographical and seasonal distribution of these ticks, particularly *R. sanguineus* in dogs, underscores the widespread nature of the problem. Factors such as the lack of acaricides treatment, traditional rural housing systems, hot and humid climates, host susceptibility, and grazing practices contribute to the proliferation of tick populations. The implications of these findings are critical for animal and human health, as ticks are vectors for numerous diseases. Effective tick control measures are urgently needed to mitigate the risks associated with tick-borne illnesses. However, the study has limitations, including its reliance on farmer interviews and the need for more comprehensive data on tick distribution and host interactions. Future work should focus on longitudinal studies to monitor tick populations, evaluate the efficacy of acaricides treatments, and explore integrated pest management strategies. Additionally, public awareness campaigns and improved veterinary practices are essential to combat tick infestations effectively. In conclusion, the fight against ticks and the diseases they transmit must remain a priority. Addressing the factors contributing to tick proliferation and implementing targeted control measures will be crucial in safeguarding animal and human health in the region. Continued research, community engagement, and policy support are vital to achieving sustainable solutions to this persistent problem.

## Consent for publication

Not applicable.

## Availability of data and materials

All data generated or analyzed during this study are included in this published article and its supplementary information files. The rest of datasets used and analyzed during the current study are available from the corresponding author on reasonable request.

## Ethics and Consent to Participate declarations

Not applicable

